# COMPASS enables cohort-independent digital biomarker discovery and pathway quantification

**DOI:** 10.64898/2025.12.02.687315

**Authors:** Saptarshi Sinha, Pradipta Ghosh

**Author notes:** Correspondence to (SS) or (P.G).

## Abstract

Reproducible and clinically transferable quantification of pathway activity remains a major barrier in precision medicine, where biomarker performance often depends on cohort composition and normalization strategies. Here, we introduce COMPASS (<u>COMP</u>osite <u>A</u>ctivity <u>S</u>coring <u>S</u>ystem), a deterministic threshold-based framework that converts gene expression into quantitative pathway activity scores without reliance on reference cohorts. COMPASS derives gene-specific activation thresholds directly from data, standardizes deviations from thresholds, and integrates directionally opposing genes into a single composite score. This enabled transparent activity scoring, statistical comparisons, and survival analyses without coding. Across diverse biological and clinical datasets, COMPASS robustly quantified cellular states, benchmarked the humanness and disease relevance of new approach methodologies, and stratified outcomes. Compared to GSVA and ssGSEA, COMPASS demonstrated greater consistency across datasets and improved robustness in bootstrap analyses, particularly for bidirectional programs, including regulatory-approved sepsis gene signatures. COMPASS therefore addresses a critical unmet need for exact, interpretable, and clinically transferable biomarker discovery and outcome modeling across diverse biological and clinical settings.

## Introduction

The rapid expansion of transcriptomic profiling, artificial intelligence/machine learning (AI/ML), and large-scale biomarker discovery efforts has led to an explosion of gene signatures proposed to define cellular states, predict outcomes, and guide therapeutic decisions. Yet, translating these signatures into clinically reliable biomarkers remains challenging because most current pathway-scoring approaches quantify relative enrichment rather than absolute biological activity. Widely used enrichment-based methods (e.g., GSEA^1^, GSVA^2^, ssGSEA^3^, PLAGE^4^, AUCell^5^, and VAM^6^) rely on evolving ontologies, cohort-dependent normalization, and permutation-based statistics, introducing variability across datasets, platforms, and populations and limiting reproducibility, interpretability, and clinical transferability ^7–9^ (**Fig. 1a***-left*). These limitations become particularly important for bidirectional gene programs, where simultaneous induction and repression of distinct genes encode biologically meaningful states that may be obscured when treated as undirected gene sets.

**Figure 1.**
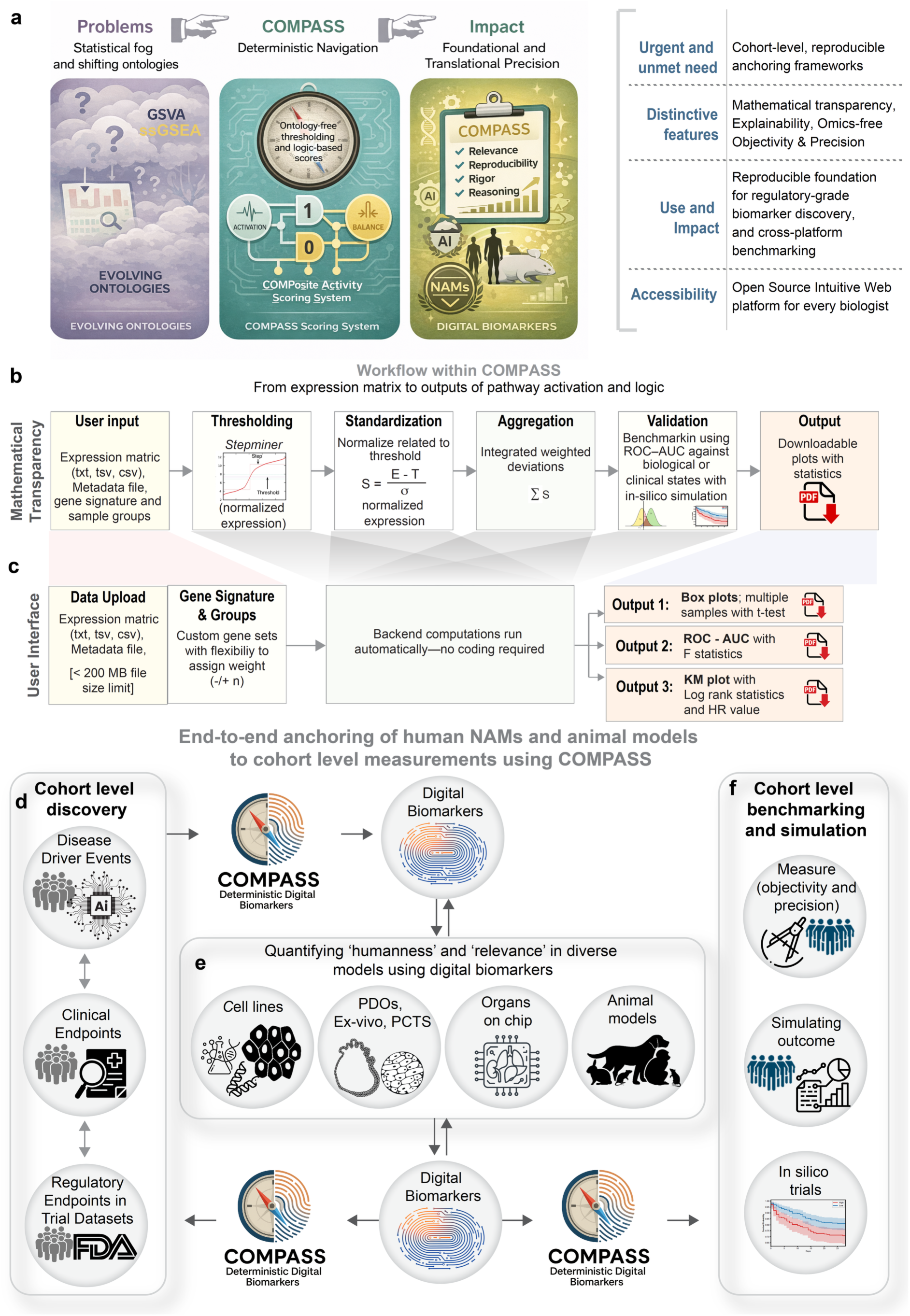
Overview of the COMPASS Framework for translational Integration of Digital Biomarkers Across Models and Cohorts Using COMPASS. The conceptual foundation (a), computational workflow (b), and workflow for translational integration of digital biomarkers for analyzing gene expression data to derive logic-based pathway activation scores and digital biomarkers. **(a) Conceptual Foundation**: COMPASS addresses limitations of traditional logic modeling by offering a deterministic, reproducible, and accessible framework for translational precision medicine. **(b) Computational Workflow**: Workflow schematic showing user inputs (expression matrix, gene signature, logic), automated backend computations, and outputs including box plots, ROC-AUC analyses and survival analyses. **(c) User Interface:** Example of an intuitive interface and downloadable outputs demonstrating COMPASS’s ability to stratify groups and assess pathway activation with statistical significance, while remaining accessible to all biologists. **(d-f) Translational Workflow:** Workflow schematic illustrates a three-part framework for aligning diverse biological models with cohort-level measurements through digital biomarkers and the COMPASS system. (**d**) **Cohort-Level Discovery**: Disease driver events, clinical endpoints, and regulatory endpoints are identified from large-n human trial datasets. These serve as anchors for downstream modeling and validation. (**e**) **Model-Level Quantification**: Diverse experimental platforms—including cell lines, patient-derived organoids (PDOs), ex vivo tissues (e.g., precision-cut tissue slices, live or fixed), organs-on-chip, and animal models—are evaluated using digital biomarkers. COMPASS objectively quantifies the humanness and relevance of each model’s response, enabling cross-comparison and translational alignment. (**f**) **Cohort-Level Benchmarking and Simulation**: Validated digital biomarkers are used to simulate cohort-level responses with high objectivity and precision. In silico trials (with survival analysis) further extend predictive capacity, supporting regulatory decision-making and personalized medicine. Central to this framework, COMPASS acts as a translational bridge linking small-n mechanistic studies with large-n population insights and enabling iterative refinement of digital biomarkers across biological scales.

Here, we introduce **COMPASS (COMPosite Activity Scoring System)**, a deterministic, ontology-free, and web-based framework that quantifies gene-set activity directly from expression data (**Fig. 1a***-middle*). COMPASS derives gene-specific activation thresholds, standardizes deviations from these boundaries, and integrates oppositely regulated genes into a single composite score using closed-form logic, enabling stable, interpretable, and clinically transferable digital biomarkers at single-sample resolution.

## Results

### A deterministic, threshold-based framework for pathway activity quantification

COMPASS operationalizes a threshold-based view of biological systems, in which cellular states emerge from transitions (high vs low) across expression boundaries [expanded here^10^]. Modeled for long using the *<u>Bo</u>*olean *<u>N</u>*etwork *<u>E</u>*xplorer (BoNE^11^) and formalized recently as **AXIOM-Bio**^10^ (*<u>A</u>bstract e<u>X</u>pression Inference and <u>O</u>ntology-free <u>M</u>odeling for <u>Bio</u>logical Logic and Disease*), this framework for creating foundational models of disease continuum states has demonstrated reproducibility across species and diseases^10, 12^. However, BoNE’s computational complexity has limited accessibility. By embedding this logic in an accessible and intuitive web platform (https://compass.precsn.com/), COMPASS enables users to interpret raw expression data using gene sets as deterministic digital biomarkers with minimal technical overhead, bridging biological reasoning, computational rigor, and translational relevance (**Fig. 1a***-right*).

Unlike enrichment-based methods (GSEA^1^, GSVA^2^, ssGSEA^3^, PLAGE^4^, AUCell^5^, and VAM^6^) that depend on ranked gene lists and random permutations, COMPASS deterministically measures biological activity without reference to predefined ontologies. In contrast to matrix-factorization or deep-learning models such as PCA, ICA, NMF, MOFA⁺, or variational autoencoders^13, 14^), COMPASS avoids latent dimensions, retaining interpretability while eliminating stochastic variation.

Beyond classification and pathway quantification, translational applications require linking molecular measurements to clinical outcomes. Because COMPASS generates continuous, direction-aware composite scores of any gene set at the sample level, these scores can be directly used to stratify samples into high and low groups for survival analyses, including Kaplan–Meier analysis and Cox proportional hazards modeling. This enables a single digital biomarker to support classification, cross-cohort benchmarking, and outcome prediction within a unified deterministic framework. Importantly, because thresholds are derived directly from the data rather than from evolving ontologies, COMPASS outputs are reproducible over time for a given dataset and gene signature, and are transferable across platforms, tissues, cohorts, and species.

### Mathematical formulation of COMPASS: thresholding, standardization, and aggregation

The COMPASS workflow transforms a raw gene-expression matrix into deterministic, quantitative measures of pathway activity through a three-step process: (i) thresholding, (ii) standardization, and (iii) aggregation (**Fig. 1b**; see ***Methods***). Each step corresponds to a concrete, interpretable biological operation rather than an abstract statistical transformation.

Briefly, in the thresholding step, COMPASS identifies for every gene its intrinsic expression boundary based on the given sample set and its inflection point where its expression pattern transitions dramatically from “off” to “on”. This data-driven threshold, determined automatically by adaptive segmentation algorithms, pinpoints where biological probability collapses into commitment. The standardization step aligns genes of varying dynamic range onto a unified scale, transforming noisy expression profiles into interpretable distance measures that indicate how far each gene lies from its activation point. Finally, the aggregation step integrates these standardized deviations across genes within a defined set, applying direction-specific weights to capture the balance of opposing influences. The resulting composite score represents the net activation of a pathway in each sample and rank them based on the gene set.

Unlike enrichment-based methods that rely on predefined ontologies or random permutations, COMPASS derives gene-specific thresholds directly from expression data, converts deviations from these thresholds into standardized metrics, and integrates them to compute composite activity scores. All computations are closed form and deterministic: identical input produces identical output, ensuring reproducibility, auditability, and mathematical transparency. The three steps are further expanded below:

**1.** *Threshold determination (biological decision boundaries)*: At the core of COMPASS lies the *StepMiner*^15^ algorithm, an adaptive algorithm originally developed to identify sharp transcriptional transitions from noisy expression data. For each gene *g*, the expression values across samples are sorted, and a step function is fit to this ordered distribution. The transition index (*Tg*) represents the inflection point that minimizes within-group variance—corresponding to the biological boundary between low (“off”) and high (“on”) expression states.

For each gene *g* and sample *i*, COMPASS computes a normalized deviation from the threshold as:

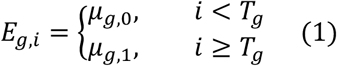

Where *T*_*g*_ denotes the transition index minimizing within-group variance. This threshold defines the binary states “low (0)” and “high (1)” expression. This adaptive segmentation identifies the intrinsic activation threshold for each gene directly from data, without requiring any external annotation or training. Conceptually, this threshold identifies where biological probability collapses into commitment (“off” → “on”)^10^. In doing so, thresholding reveals the cellular decision boundary.

To minimize stochastic fluctuations around this boundary, COMPASS applies a small confidence offset (σ) to the threshold, defining the final decision boundary of each gene *g* as:

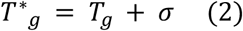

where, σ = 0.5 log_2_ *Units*. This conservative shift in threshold ensures that only values confidently above or below the step contribute to the score.

**2.** *Standardization (normalization across genes):* Once thresholds are established, each gene’s deviation from its intrinsic boundary is standardized relative to its variance, yielding a dimensionless measure of distance from the decision point: For every gene *g* and sample *i*, the deviation from this intrinsic threshold is standardized as:

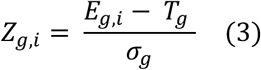

where *E_g,i_* is the observed expression and σ_*g*_ the gene-specific standard deviation. *Z_g,i_* thus measures the distance from the decision boundary^11^, producing a hybrid metric: discrete when categorical precision is needed (i.e., digital) and continuous when population gradients matter (i.e., analog). The standardized deviation used in COMPASS is then calculated as:

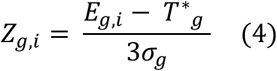

Dividing by 3σ_g_(instead of 1×σ_g_) compresses extreme outliers and brings all genes to a comparable dynamic scale. This preserves biological directionality while reducing threshold-adjacent noise.

This standardization step converts heterogeneous expression profiles into a common coordinate system in which each *Z*₍*g,i*₎ representing how strongly a gene deviates from its activation threshold.

**3.** *Composite aggregation (pathway-level integration)*: For any gene-set, *G* = {*g*_1_, *g*_2_, …, *g*_*m*_}, COMPASS computes the composite activity score for each sample *i* as the weighted mean of standardized deviations:

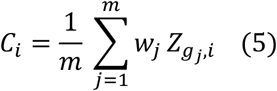

where weights *wⱼ* encode gene directionality (+1 = activation, −1 = repression). The resulting composite score (C_i_) reflects the net balance of activation and repression, providing an interpretable, quantitative index of pathway activity. A higher positive value indicates dominance of activation, while a negative value indicates repression. Published use cases include pro-versus anti-inflammatory macrophage states^16^, differentiation versus stemness^17^, or metabolic activation versus inhibition, within a single continuous framework.

This closed-form calculation eliminates the randomness inherent to permutation-based enrichment methods. It also ensures that each component—expression, threshold, deviation, and directionality—corresponds to an observable biological quantity rather than a statistical abstraction.

To evaluate biological and clinical relevance, COMPASS rank the sample set based on the derived composite scores and perform a ROC–AUC analysis, comparing the classification accuracy of defined sample groups (for example, disease vs control or responder vs non-responder) based on another *Stepminer* threshold computed on the composite scores. The Area Under the Curve (AUC) is computed as:

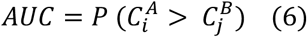

where *C^A^_i_* and *C^B^_j_* denote the composite scores of state *A* (e.g., treated or disease) and state *B* (e.g., control or healthy) samples, respectively. This provides a universal, model-agnostic measure of discriminatory power that reflects the internal biological coherence of each gene-set’s activity pattern and serves as a statistical summary the same. Because all operations are deterministic, identical inputs produce identical outputs, ensuring reproducibility and traceability across runs, datasets, and environments.

### A web-based platform for reproducible, code-free pathway activity scoring

COMPASS is implemented as a web-based analytical platform (https://compass.precsn.com/) with an intuitive graphical interface (**Fig. 1c; Figure Supplement S1**). Users upload a tab- or comma-delimited gene expression matrix (txt, tsv, or csv), optionally include time-to-event metadata, define gene signatures with directionality, and initiate analysis through a point-and-click workflow. All computations, including thresholding, standardization, aggregation, and statistical evaluation, are executed automatically in the background, without the need for pathway databases or coding. Gene-set definitions are fully customizable, with flexibility to assign directionality weights (+/−).

The platform generates publication-ready outputs, including boxplots for group comparisons (accompanied by pairwise *t*-test statistics), ROC–AUC analyses (with associated *F* and *p* values), composite activity tables (containing standardized *Z*-scores and composite scores for each pathway and sample), and survival analyses (with hazard ratios and log-rank statistics). Results are downloadable in standard formats (CSV, PDF).

In terms of accessibility, unlike conventional computational frameworks, COMPASS is designed to be intuitive and accessible to both computational and non-computational users. Users can define and compare any number of sample groups directly from their uploaded data (e.g., WT vs KO, responders vs non-responders, multiple treatment conditions). The interface enables point-and-click selection of genes, pathways, and groups, generating publication-ready figures without manual scripting (**Fig. S1a-b**). All statistical analyses, such as composite score calculation, group-wise comparisons, ROC–AUC, survival analyses are performed automatically. By integrating accessibility with mathematical transparency, COMPASS enables consistent, sample-level quantification of pathway activity across biological and clinical contexts.

### Clinically anchored benchmarking across multi-cohort datasets

COMPASS supports diverse use cases (see **Fig. 1d-f**): benchmarking organoids, animal models, or New Approach Methodologies (**NAMs**) against human cohorts to quantify *humanness* and biological *relevance*^17–23^; computing composite activity scores predictive of outcomes^17^; tracking drug response or toxicity^21^; and mapping disease-driver cell states such as alveolar cytopathy, immune cell processes (reactivity vs. tolerance) or differentiation^11, 16, 17, 24^.

Across diverse gene signatures and datasets spanning oncology, infectious disease, inflammatory disorders, and cellular state mapping (**Fig. 2a-o**), COMPASS consistently recapitulated expected biological patterns across human cohorts, animal models, and organoid systems. In colorectal cancer, a stemness signature quantified concordant therapeutic responses across cell lines, xenografts, and patient-derived organoids. In colorectal cancer, a 50-gene stemness signature^17^ quantified the efficacy of a differentiation-inducing therapeutic across cell lines, xenografts, and patient-derived organoids, revealing concordant pathway suppression in all models (**Fig. 2a–c**). In respiratory viral injury^25^, COMPASS detected induction of a lung damage signature across infected human biopsies, hamster lungs, and adult stem cell–derived lung organoids (**Fig. 2d–f**), highlighting conserved molecular pathology.

**Figure 2.**
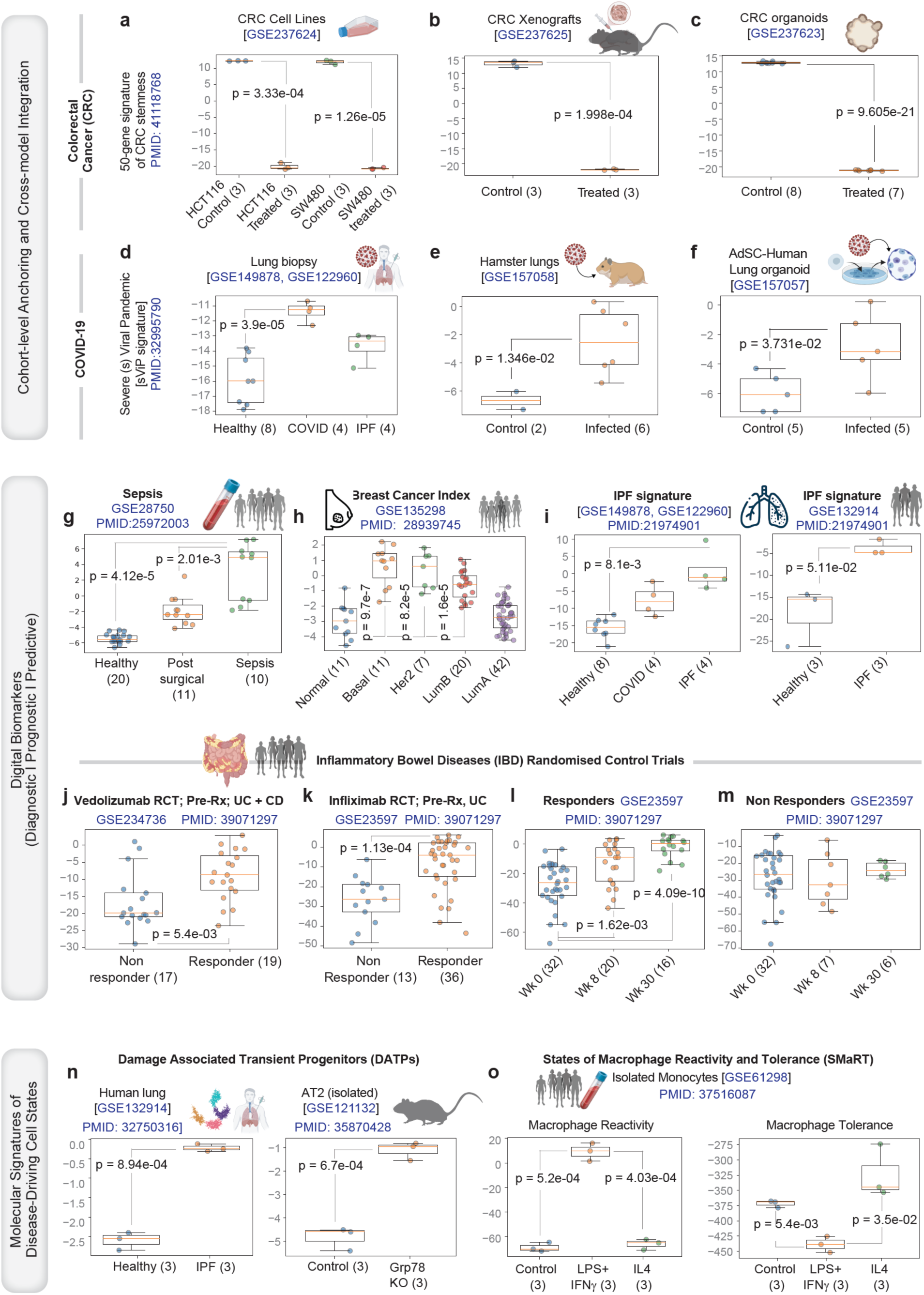
Cohort-level anchoring and cross-model benchmarking using COMPASS. COMPASS enables deterministic quantification of pathway activity across cohorts, tissues, species, and disease models, allowing unified navigation of biological logic and clinical relevance. Published examples illustrating such cross-species and cross-model ontology-free navigation between human, organoid, and animal systems are shown. **(a–c)** *Colorectal cancer (CRC)*: A 50-gene CRC stemness signature^17^ was used to score the efficacy of a first-in-class differentiation inducing therapeutic agent across cell lines (a), xenografts (b), and patient-derived organoids (c), demonstrating consistent efficacy across experimental models^17^. **(d–f)** *COVID-19 and lung injury*: A 20-gene signature^25^ indicative of severe alveolar damage in the setting of respiratory viral pandemics is induced in CoV-infected human lungs^33^ (d), hamster lungs^34^ (e), and human adult stem cell-derived lung organoids^23^ (f), recapitulating conserved pathway activation across species and model systems (datasets and PMIDs annotated). **(g–i)** *Diagnostic and prognostic digital biomarkers*: Distinct gene sets corresponding to sepsis (g; 11-gene diagnostic gene set, identified using a comprehensive time-course-based multicohort analysis of sepsis^26^), breast cancer subtypes (h; 7-gene **BCI**™ signature, the only genomic test that is both prognostic and predictive and is recognized by the NCCN and the ASCO^®^ Clinical Practice Guideline to identify patients who might benefit from adjuvant anti-estrogen therapy beyond 5 years), and idiopathic pulmonary fibrosis (I; a 153-gene IPF diagnostic signature^28^, comprised of 76 up and 77 down genes) generated robust discriminatory activity scores, matching known clinical groupings. **(j–m)** *Digital biomarkers for therapeutic response prediction and tracking*: A 34-gene signature^21^, indicative of mucosal barrier bioenergetic state, predicts therapeutic response in Inflammatory bowel disease (IBD) clinical trial datasets, distinguishing responders (high-34) from non-responders (low-34), regardless of class of drug (j-k) and tracks remission during therapy among responders in longitudinal datasets. Wk, week; UC, Ulcerative colitis; CD, Crohn’s disease. **(n–o)** *Molecular signatures of disease-driving cell states*: A 6-gene signature of damage-associated transient progenitors (DATP^29^) is induced in human IPF and murine models that recapitulate fundamental processes in IPF (i.e., impaired ER stress response^34^; n). Gene signatures of macrophage^16^ reactivity (48 genes) and tolerance (232 genes) accurately captured macrophage polarization continua (reactivity vs tolerance; o) across cytokine-stimulated monocytes, revealing deterministic transitions that define physiologic and pathologic spectra.

Clinical (diagnostic and prognostic) signatures for sepsis^26^, breast cancer^27^, and idiopathic pulmonary fibrosis^28^ generated robust discriminatory scores aligned with clinical classifications (**Fig. 2g–i**). In two independent IBD clinical trials, a bioenergetic barrier signature^21^ stratified responders from non-responders across therapeutic classes and tracked remission longitudinally (**Fig. 2j–m**). Finally, COMPASS quantified cell-state transitions—capturing induction of DATPs in fibrosis models^29^ and defining graded polarization from macrophage reactivity to tolerance^16^ (**Fig. 2n–o**).

These results demonstrate consistent performance across cohorts, models, and species, supporting the use of COMPASS for cross-context benchmarking and outcome-linked digital biomarker modeling.

### Benchmarking against other methodologies: Reduced variability and enhanced robustness under resampling

**Table 1** highlights key differences between COMPASS and other enrichment-based gene set scoring methods. Unlike other approaches, which compute relative, cohort-dependent scores, COMPASS derives gene-specific thresholds and generates absolute, direction-aware activity scores at the sample level. This deterministic formulation avoids permutations, improves reproducibility, and enables stable integration of opposing gene signals. In addition, COMPASS supports multi-class and continuous analyses, facilitating consistent pathway quantification across datasets and biological contexts.

**Table 1.**
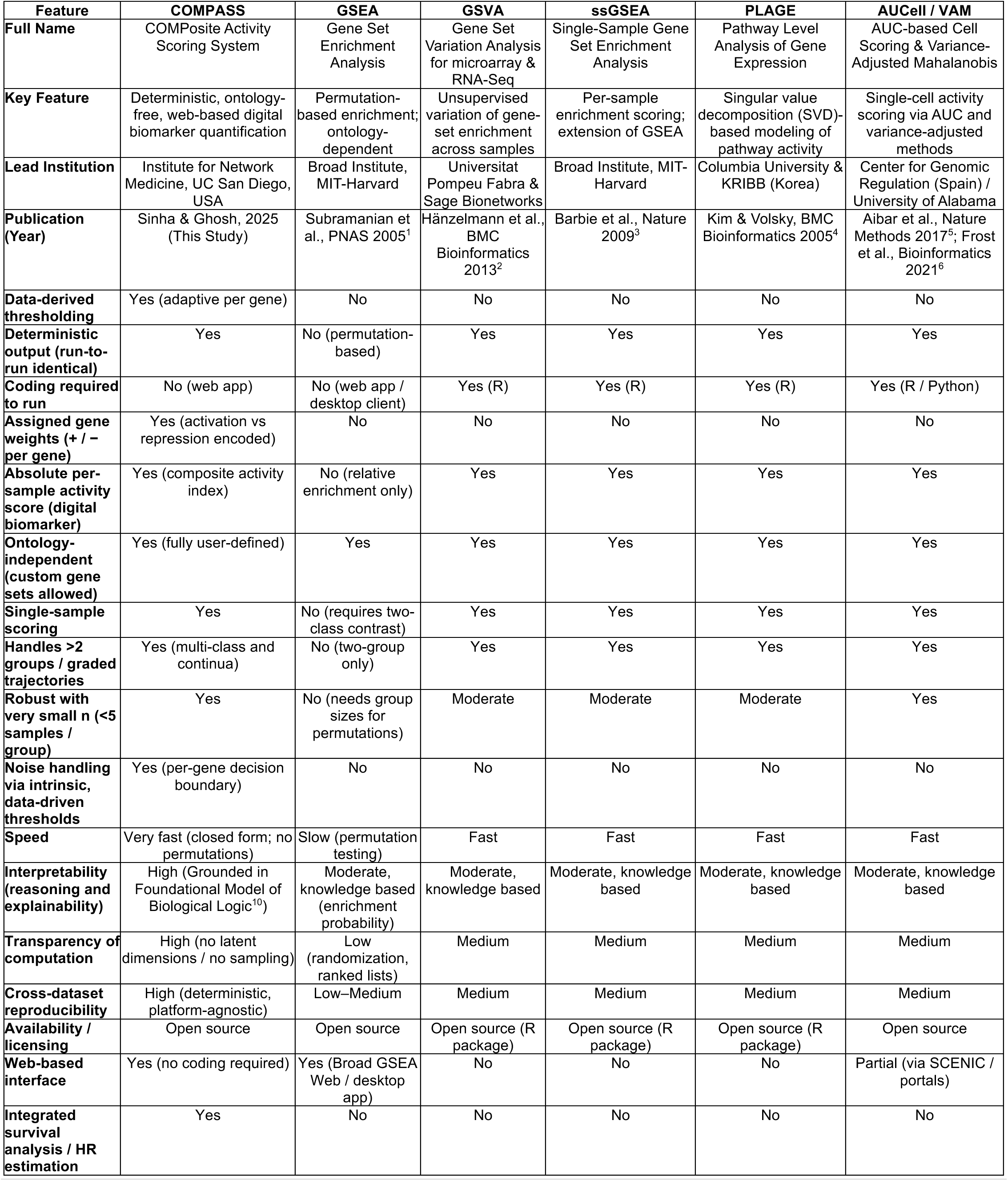
Comparative features of COMPASS versus existing gene-set activity frameworks. Acronyms: GSEA, Gene Set Enrichment Analysis; GSVA, Gene Set Variation Analysis; ssGSEA, Single-Sample Gene Set Enrichment Analysis; PLAGE, Pathway-Level Analysis of Gene Expression; AUCell, Area-Under-the-Curve Cell Scoring; VAM, Variance-Adjusted Mahalanobis.

| Feature | COMPASS | GSEA | GSVA | ssGSEA | PLAGE | AUCCell / VAM |
| --- | --- | --- | --- | --- | --- | --- |
| **Full Name** | COMPosite Activity Scoring System | Gene Set Enrichment Analysis | Gene Set Variation Analysis for microarray & RNA-Seq | Single-Sample Gene Set Enrichment Analysis | Pathway Level Analysis of Gene Expression | AUC-based Cell Scoring & Variance-Adjusted Mahalanobis |
| **Key Feature** | Deterministic, ontology-free, web-based digital biomarker quantification | Permutation-based enrichment; ontology-dependent | Unsupervised variation of gene-set enrichment across samples | Per-sample enrichment scoring; extension of GSEA | Singular value decomposition (SVD)-based modeling of pathway activity | Single-cell activity scoring via AUC and variance-adjusted methods |
| **Lead Institution** | Institute for Network Medicine, UC San Diego, USA | Broad Institute, MIT-Harvard | Universitat Pompeu Fabra & Sage Bionetworks | Broad Institute, MIT-Harvard | Columbia University & KRIBB (Korea) | Center for Genomic Regulation (Spain) / University of Alabama |
| **Publication (Year)** | Sinha & Ghosh, 2025 (This Study) | Subramanian et al., PNAS 2005<sup>1</sup> | Hänzelmann et al., BMC Bioinformatics 2013<sup>2</sup> | Barbie et al., Nature 2009<sup>3</sup> | Kim & Volsky, BMC Bioinformatics 2005<sup>4</sup> | Aibar et al., Nature Methods 2017<sup>5</sup>; Frost et al., Bioinformatics 2021<sup>6</sup> |
| **Data-derived thresholding** | Yes (adaptive per gene) | No | No | No | No | No |
| **Deterministic output (run-to-run identical)** | Yes | No (permutation-based) | Yes | Yes | Yes | Yes |
| **Coding required to run** | No (web app) | No (web app / desktop client) | Yes (R) | Yes (R) | Yes (R) | Yes (R / Python) |
| **Assigned gene weights (+ / - per gene)** | Yes (activation vs repression encoded) | No | No | No | No | No |
| **Absolute per-sample activity score (digital biomarker)** | Yes (composite activity index) | No (relative enrichment only) | Yes | Yes | Yes | Yes |
| **Ontology-independent (custom gene sets allowed)** | Yes (fully user-defined) | Yes | Yes | Yes | Yes | Yes |
| **Single-sample scoring** | Yes | No (requires two-class contrast) | Yes | Yes | Yes | Yes |
| **Handles >2 groups / graded trajectories** | Yes (multi-class and continua) | No (two-group only) | Yes | Yes | Yes | Yes |
| **Robust with very small n (<5 samples / group)** | Yes | No (needs group sizes for permutations) | Moderate | Moderate | Moderate | Yes |
| **Noise handling via intrinsic, data-driven thresholds** | Yes (per-gene decision boundary) | No | No | No | No | No |
| **Speed** | Very fast (closed form; no permutations) | Slow (permutation testing) | Fast | Fast | Fast | Fast |
| **Interpretability (reasoning and explainability)** | High (Grounded in Foundational Model of Biological Logic<sup>10</sup>) | Moderate, knowledge based (enrichment probability) | Moderate, knowledge based | Moderate, knowledge based | Moderate, knowledge based | Moderate, knowledge based |
| **Transparency of computation** | High (no latent dimensions / no sampling) | Low (randomization, ranked lists) | Medium | Medium | Medium | Medium |
| **Cross-dataset reproducibility** | High (deterministic, platform-agnostic) | Low-Medium | Medium | Medium | Medium | Medium |
| **Availability / licensing** | Open source | Open source | Open source (R package) | Open source (R package) | Open source (R package) | Open source |
| **Web-based interface** | Yes (no coding required) | Yes (Broad GSEA Web / desktop app) | No | No | No | Partial (via SCENIC / portals) |
| **Integrated survival analysis / HR estimation** | Yes | No | No | No | No | No |

To position COMPASS relative to widely used single-sample scoring methods, we performed head-to-head comparisons against GSVA and ssGSEA across 10 independent cohorts (comprised of 232 control and 499 sepsis samples) spanning diverse clinical contexts and patient populations using a clinically validated 11-gene sepsis signature^26^. This signature was originally identified and rigorously validated in multi-cohort datasets^26, 30^ (**Fig. 3a**), and recently approved by the FDA for marketing. To assess the role of gene directionality, analyses were conducted using upregulated genes, downregulated genes, and the full composite signature.

**Figure 3.**
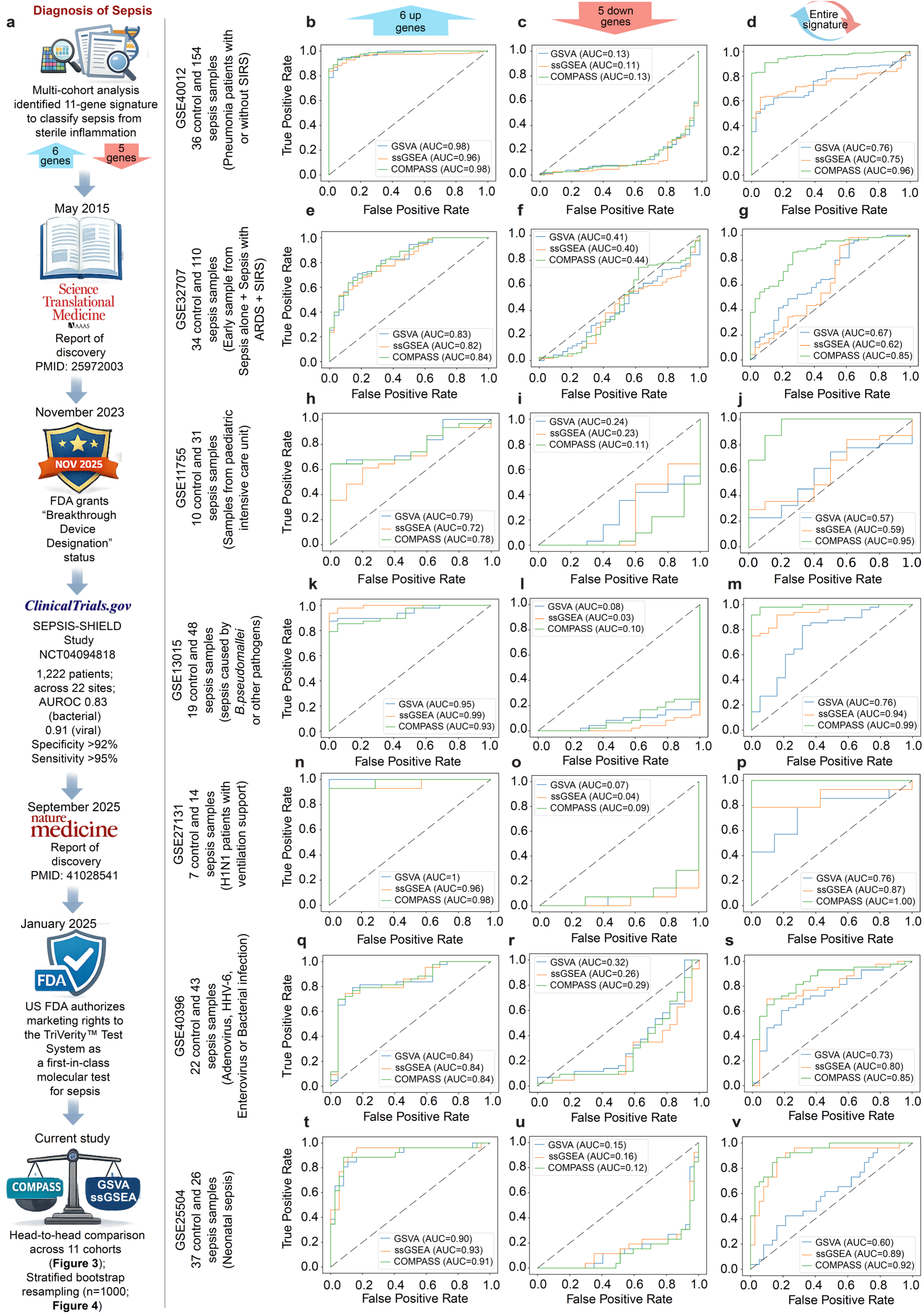
Clinically anchored benchmarking of COMPASS across multi-cohort sepsis datasets. **(a)** Schematic represents the clinical and translational trajectory of the 11-gene host-response signature for sepsis. From left to right: A multi-cohort discovery framework identified a minimal gene set that distinguishes sepsis from sterile inflammation, followed by large-scale validation across independent, multi-center cohorts (including SEPSIS-SHIELD^30^), demonstrating high diagnostic performance and enabling regulatory advancement to an FDA-authorized molecular diagnostic. This clinically validated gene signature provides a digital gene signature and an opportunity for head-to-head comparison of COMPASS with existing analyses frameworks GSVA and ssGSEA for their ability to accurately and reproducibly classify clinically determined sepsis samples from healthy controls across 11 diverse cohorts. (**b-v**) Receiver operating characteristic (ROC) curves comparing GSVA (blue), ssGSEA (orange), and COMPASS (green) across independent transcriptomic cohorts. Panels are arranged by gene set composition (columns) and cohort (rows). Columns represent: left, 6 upregulated genes; middle, 5 downregulated genes; right, full 11-gene signature. Rows correspond to distinct clinical cohorts spanning discovery and validation phases: (**b-d**) <u>GSE40012</u> (pneumonia patients with or without Systemic Inflammatory Response Syndrome [SIRS]), (**e-g**) <u>GSE32707</u> (early sepsis, SIRS, and acute respiratory distress syndrome [ARDS] samples), (**h-j**) <u>GSE11755</u> (pediatric intensive care unit [ICU] cohort), (**k-m**) <u>GSE13015</u> (pathogen-driven sepsis), (**n-p**) <u>GSE27131</u> (H1N1 with respiratory failure), (**q-s**) <u>GSE40396</u> (viral and bacterial infections), (**t-v**) <u>GSE25504</u> (neonatal sepsis). Across cohorts, enrichment-based methods (GSVA, ssGSEA) show context-dependent performance, particularly when gene directionality is partitioned (up vs down genes). In contrast, COMPASS demonstrates robust and consistent discrimination, with maximal performance when applied to the full bidirectional signature, highlighting its ability to integrate opposing gene programs into a unified, outcome-relevant score. Additional cohorts are shown in **Figure S2**.

Across cohorts, GSVA and ssGSEA showed variable performance depending on gene subset and cohort composition, particularly when directional (upregulated or downregulated) components were evaluated separately (for example, **Fig. 3b-c**; **Figure Supplement S2a-b**). By contrast, COMPASS maintained consistent discrimination when integrating directionally opposing genes into a single composite score (for example, **Fig. 3d**; **Figure Supplement S2c**), highlighting the importance of direction-aware aggregation for stable pathway quantification.

Robustness was evaluated using stratified bootstrap resampling (n = 1000; **Fig. 4a**). For each bootstrap replicate, AUC was recalculated, with score direction aligned when necessary to ensure consistent interpretation. Sensitivity and specificity were estimated using a fixed decision threshold for each method, determined once from the original dataset by maximizing the Youden index and then applied uniformly across all bootstrap replicates. COMPASS exhibited tighter AUC distributions and higher central tendency across cohorts (**Fig. 4b-k**), indicating reduced variance and stability under sampling perturbation. GSVA and ssGSEA showed broader distributions, reflecting greater sensitivity to cohort composition/resampling. Quantitative summaries (**Table 2**) further support these findings. COMPASS achieved balanced performance across cohorts, (mean specificity of 0.92 and sensitivity of 0.91). Notably, these values approach the performance range reported for the SEPSIS-SHIELD clinical study^30^ (specificity >92% and sensitivity >95%), indicating that COMPASS-derived scores recapitulate clinically relevant discriminatory power at the cohort level. By contrast, GSVA (specificity 0.69, sensitivity 0.69) and ssGSEA (specificity 0.80, sensitivity 0.88) showed substantially lower and less balanced performance.

**Figure 4.**
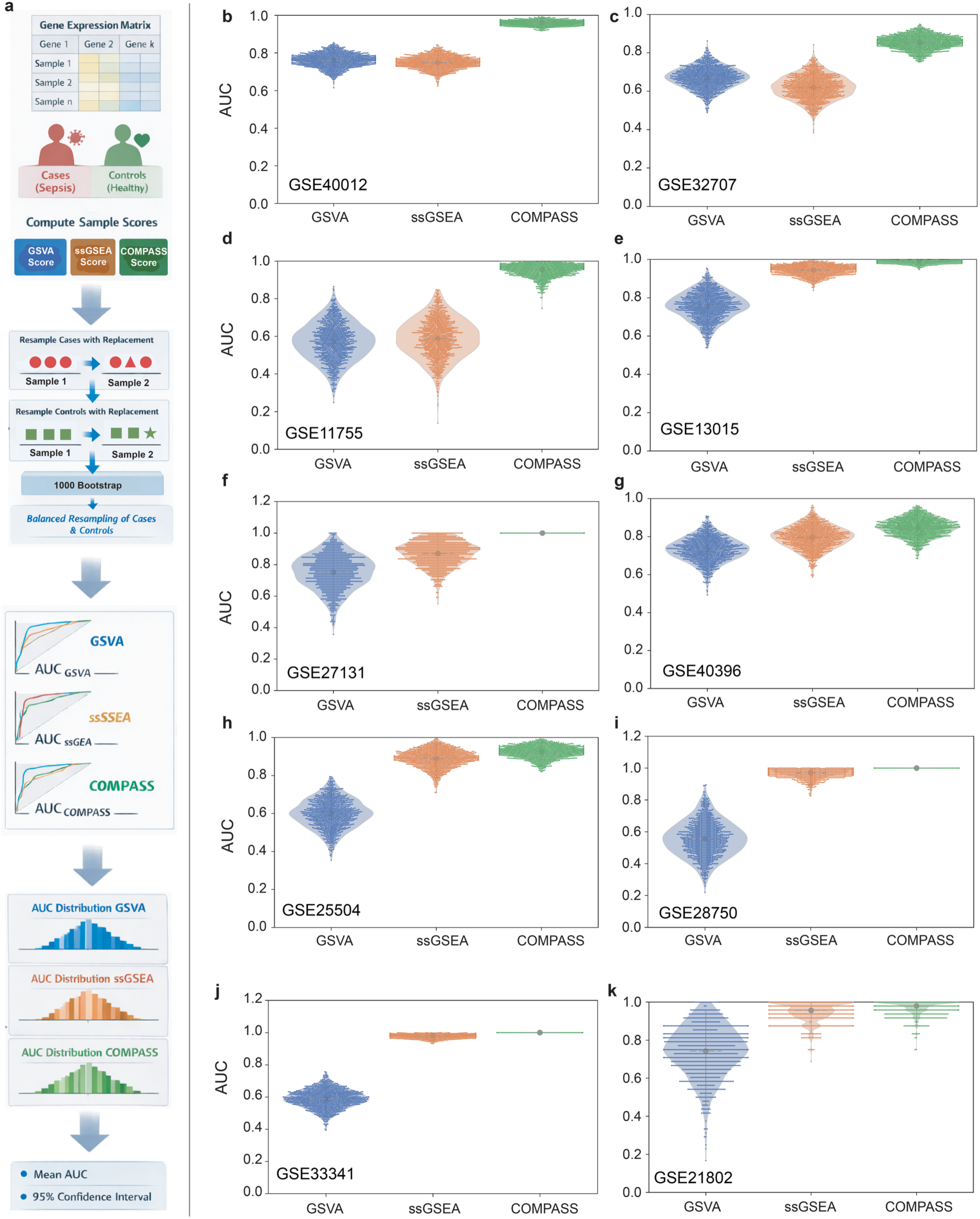
Comparative robustness of GSVA, ssGSEA, and COMPASS assessed by stratified bootstrap resampling across independent cohorts. **(a)** Schematic of the bootstrap workflow (top to bottom). Gene expression matrices are used to compute sample-level scores for GSVA, ssGSEA, and COMPASS. Samples are then resampled with replacement within each class (cases and controls) to preserve class balance. This procedure is repeated for 1000 iterations. In each iteration, score direction is aligned when necessary to ensure consistent interpretation, and receiver operating characteristic (ROC) curves are generated to compute area under the curve (AUC). The resulting AUC distributions are summarized to estimate variability and confidence intervals for each method (see **Table 2**). (**b–k**) Violin plots showing distributions of AUC values across 1000 bootstrap iterations for GSVA (blue), ssGSEA (orange), and COMPASS (green) in independent cohorts. Each panel corresponds to a dataset used in Figure 3 and **Figure S2**. Each violin represents the distribution of resampled AUC values under class-balanced conditions.

**Table 2.** Stratified bootstrap performance of GSVA, ssGSEA, and COMPASS across independent cohorts: Performance metrics are summarized from bootstrap resampling (n = 1000) performed independently within each class (cases and controls) to preserve class balance. For each iteration, receiver operating characteristic (ROC) curves were used to derive area under the curve (AUC) values, from which mean AUC and 95% confidence intervals (CI) were estimated. Sensitivity and specificity were calculated using a fixed decision threshold for each method, determined from the original dataset by maximizing the Youden index and applied consistently across iterations. Sensitivity and specificity are presented as mean ± standard deviation (S.D) in each cohort (left), and as averages across bootstrap iterations (right).

| Dataset | Method | Mean AUC | 95% CI (AUC) | Sensitivity (mean ± SD) | Specificity (mean ± SD) |
| --- | --- | --- | --- | --- | --- |
| GSE40012 | GSVA | 0.760 | (0.683-0.828) | 0.624 ± 0.038 | 0.861 ± 0.058 |
| | ssGSEA | 0.749 | (0.683-0.812) | 0.631 ± 0.037 | 0.946 ± 0.038 |
|  | **COMPASS** | **0.960** | **(0.935-0.983)** | **0.910 ± 0.023** | **0.917 ± 0.048** |
| GSE32707 | GSVA | 0.670 | (0.563-0.773) | 0.935 ± 0.024 | 0.376 ± 0.084 |
| | ssGSEA | 0.619 | (0.497-0.735) | 0.982 ± 0.012 | 0.376 ± 0.084 |
|  | **COMPASS** | **0.854** | **(0.780-0.921)** | **0.864 ± 0.033** | **0.736 ± 0.077** |
| GSE11755 | GSVA | 0.571 | (0.377-0.755) | 0.743 ± 0.079 | 0.494 ± 0.158 |
| | ssGSEA | 0.589 | (0.390-0.784) | 0.292 ± 0.079 | 1.000 ± 0.000 |
|  | **COMPASS** | **0.955** | **(0.865-1.000)** | **1.000 ± 0.000** | **0.801 ± 0.126** |
| GSE13015 | GSVA | 0.757 | (0.612-0.882) | 0.836 ± 0.053 | 0.685 ± 0.109 |
| | ssGSEA | 0.945 | (0.888-0.986) | 0.918 ± 0.040 | 0.843 ± 0.085 |
|  | **COMPASS** | **0.990** | **(0.969-1.000)** | **0.980 ± 0.020** | **0.947 ± 0.052** |
| GSE27131 | GSVA | 0.753 | (0.520-0.939) | 0.789 ± 0.107 | 0.712 ± 0.172 |
| | ssGSEA | 0.871 | (0.704-1.000) | 0.790 ± 0.106 | 1.000 ± 0.000 |
|  | **COMPASS** | **1.000** | **(1.000-1.000)** | **1.000 ± 0.000** | **1.000 ± 0.000** |
| GSE40396 | GSVA | 0.731 | (0.600-0.851) | 0.605 ± 0.078 | 0.820 ± 0.081 |
| | ssGSEA | 0.795 | (0.675-0.903) | 0.696 ± 0.070 | 0.908 ± 0.062 |
|  | **COMPASS** | **0.847** | **(0.749-0.936)** | **0.698 ± 0.071** | **0.864 ± 0.071** |
| GSE25504 | GSVA | 0.599 | (0.446-0.737) | 0.423 ± 0.097 | 0.787 ± 0.068 |
| | ssGSEA | 0.889 | (0.788-0.967) | 0.884 ± 0.062 | 0.837 ± 0.062 |
|  | **COMPASS** | **0.924** | **(0.850-0.976)** | **0.885 ± 0.062** | **0.835 ± 0.060** |
| GSE28750 | GSVA | 0.556 | (0.345-0.780) | 0.896 ± 0.096 | 0.548 ± 0.112 |
| | ssGSEA | 0.970 | (0.895-1.000) | 0.901 ± 0.096 | 0.949 ± 0.049 |
|  | **COMPASS** | **1.000** | **(1.000-1.000)** | **1.000 ± 0.000** | **1.000 ± 0.000** |
| GSE33341 | GSVA | 0.592 | (0.507-0.695) | 0.575 ± 0.248 | 0.653 ± 0.245 |
| | ssGSEA | 0.980 | (0.953-0.997) | 0.946 ± 0.033 | 0.967 ± 0.030 |
|  | **COMPASS** | **1.000** | **(1.000-1.000)** | **1.000 ± 0.000** | **1.000 ± 0.000** |
| GSE21802 | GSVA | 0.742 | (0.458-0.958) | 0.497 ± 0.141 | 1.000 ± 0.000 |
| | ssGSEA | 0.956 | (0.833-1.000) | 0.914 ± 0.081 | 1.000 ± 0.000 |
|  | **COMPASS** | **0.979** | **(0.875-1.000)** | **0.916 ± 0.082** | **1.000 ± 0.000** |

Together, these analyses demonstrate that direction-aware, deterministic scoring improves stability and reproducibility of pathway activity quantification across cohorts.

### Outcome prediction and survival stratification using COMPASS-derived scores

To assess clinical relevance, we analyzed a large multi-cohort sepsis dataset (**GSE310929**; n = 593 patients) with time to death annotation using a validated 12-gene sepsis-severity signature^30, 31^ (**Fig. 5a-b**). As in the case of the 11-gene diagnostic signature we used earlier (**Fig. 3a**), this 12-gene severity signature was also rigorously validated in the SEPSIS-SHIELD clinical study^30^ before receiving regulatory approval for marketing. Enrichment-based methods, as well as COMPASS showed comparable performance in sample classification (**Fig. 5c-e**); all underperformed to similar extent. Importantly, COMPASS-derived scores enabled direct stratification of patients by survival outcome (**Fig. 5f**). Patients with higher composite scores exhibited significantly worse survival (Hazard ratio [H.R] = 2.32), demonstrating that deterministic, direction-aware activity scores can be extended from group discrimination to time-to-event analysis.

**Figure 5.**
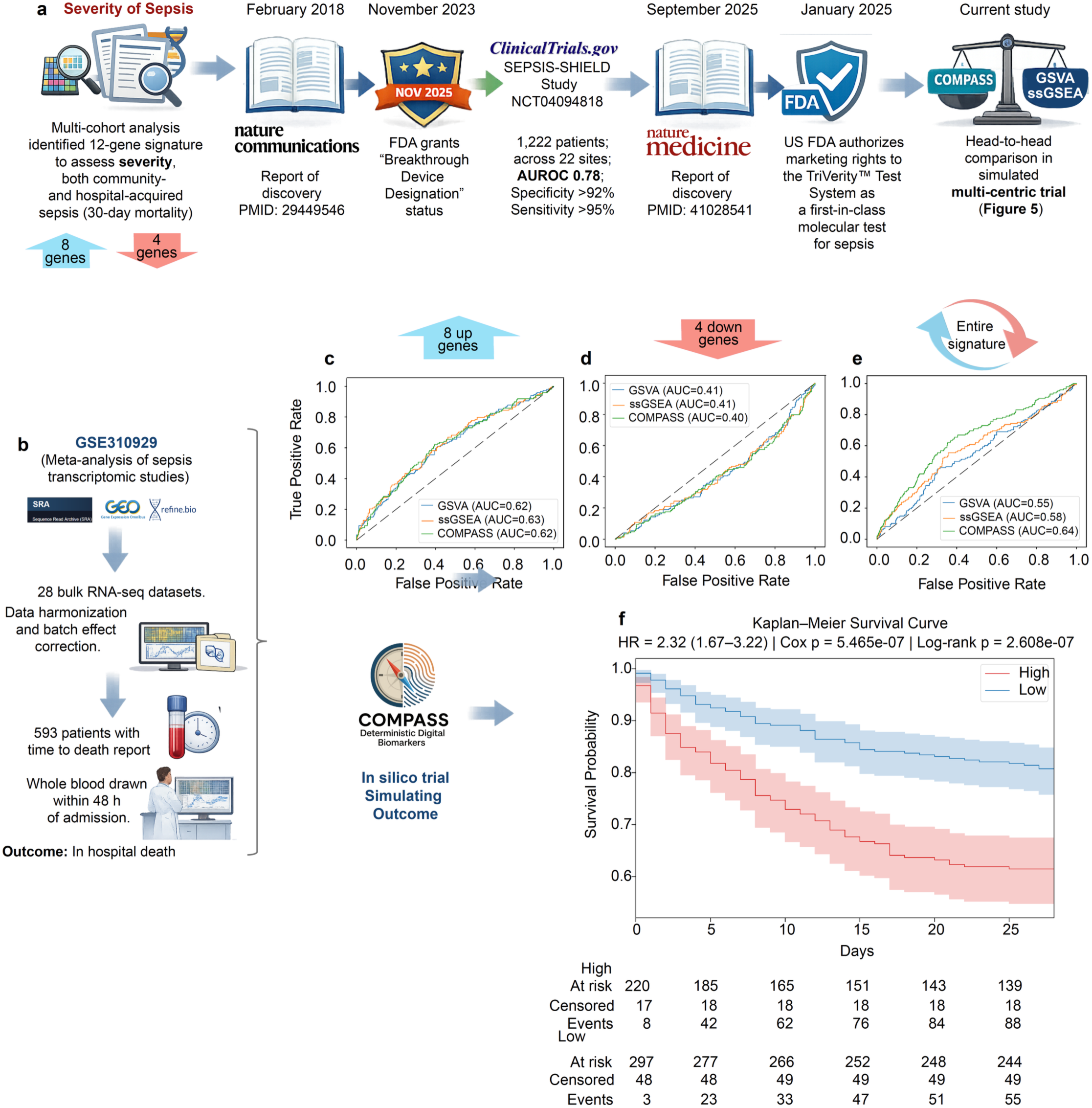
Outcome prediction and survival stratification using a clinically validated sepsis severity signature. **(a)** Clinical and translational trajectory of the 12-gene sepsis severity signature (Stanford mortality score^31^). A multi-cohort analysis identified^31^ a gene signature associated with sepsis severity and 30-day mortality across community- and hospital-acquired infections. Subsequent validation^30^ across multi-center studies, including the SEPSIS-SHIELD study^30^, demonstrated clinical utility and supported regulatory advancement, culminating in FDA authorization of a first-in-class molecular diagnostic test. In the current study, this signature is used for head-to-head comparison and outcome modeling. **(b)** Composition of the meta-analysis cohort (<u>GSE310929</u>). Expression data from 28 bulk RNA-seq datasets were harmonized and batch-corrected, yielding 593 patients with time-to-event annotations. Whole blood samples were collected within 48 hours of admission, with in-hospital mortality as the primary outcome. (**c–e**) Receiver operating characteristic (ROC) curves comparing GSVA (blue), ssGSEA (orange), and COMPASS (green) for outcome prediction across different gene set compositions. (**C**) 8 upregulated genes (**D**) 4 downregulated genes (**E**) full 12-gene signature. Each panel shows ROC curves for GSVA (blue), ssGSEA (orange), and COMPASS (green), with corresponding area under the curve (AUC) values. (**f**) Kaplan–Meier survival analysis based on COMPASS-derived scores using the full 12-gene signature. Patients stratified into high versus low score groups show significantly different survival probabilities, with worse outcomes in the high-score group (hazard ratio [HR] = 2.32, 95% CI: 1.67–3.22; Cox and log rank p-value and log-rank p-values as indicated). Shaded regions represent confidence intervals, and numbers at risk are shown below the plot.

These results establish that COMPASS provides a unified framework in which pathway activity scores function as outcome-linked digital biomarkers, enabling consistent transition from classification to survival modeling within the same analytical system.

## Discussion

Translation frequently fails when biological signals identified in discovery cohorts cannot be reproducibly anchored across validation cohorts, preclinical model systems, and patient populations. COMPASS was designed to address this gap. By explicitly modeling gene directionality and anchoring expression to intrinsic activation thresholds, COMPASS provides a deterministic framework that improves interpretability, reduces cohort dependency, and preserves biological meaning across datasets. The benchmarking analyses presented here position COMPASS alongside existing single-sample scoring methods rather than replacing them. Methods such as GSVA and ssGSEA remain valuable for relative enrichment analysis. However, as AI/ML-driven biomarker discovery accelerates and transcriptomic signatures proliferate across biomedical research, a growing challenge is ensuring that molecular measurements remain stable, allow multi-group comparison, reproducible, interpretable, and transferable across cohorts, platforms, and clinical settings. By explicitly modeling gene directionality and anchoring expression to intrinsic thresholds, COMPASS provides a deterministic framework that improves interpretability and stability. Beyond that, a conceptual advance that is operationalized through COMPASS, reduces cohort dependency, and preserves biological meaning across datasets.

A key conceptual advance of COMPASS is its ability to integrate opposing biological signals within composite signatures. This is particularly important for clinically relevant programs in which simultaneous induction and repression of distinct genes encode disease state, therapeutic response, or risk. Bootstrap analyses demonstrated reduced variability and greater consistency across cohorts, including superiority for bidirectional programs and regulatory-approved sepsis gene signatures.

Another key advance of COMPASS is its ability to directly connect pathway activity with clinically meaningful outcomes. By generating continuous, direction-aware scores at single-sample resolution, COMPASS enables objective evaluation of biomarkers and therapeutic perturbations across independent cohorts before prospective testing. The same digital biomarker can support classification, benchmarking, longitudinal monitoring, survival analyses, and outcome modeling without stochastic inference or intermediate model fitting. Such portability and reproducibility are increasingly critical as transcriptomic biomarkers move toward regulatory qualification, decentralized testing, and AI-enabled clinical decision support.

More broadly, COMPASS establishes a generalizable framework for quantitative biomarker science. Its deterministic design ensures that identical inputs yield identical outputs, enabling reproducibility across datasets, platforms, and settings, including regulatory contexts. By linking expression, threshold, deviation, and directionality within a transparent, closed-form system, COMPASS converts gene-expression data into interpretable and transferable measures of biological state, providing a foundation for consistent and scalable analysis across model systems and clinical cohorts.

This study has limitations. Because COMPASS derives thresholds directly from input data, its performance depends on data quality and preprocessing (e.g., normalization) and may be affected by limited dynamic range or residual batch effects. The framework also relies on biologically informed gene signatures with correctly assigned directionality. In its current form, COMPASS aggregates genes linearly and does not explicitly model higher-order network interactions. Additional validation across multimodal datasets, single-cell and spatial platforms, and prospective clinical studies will be important to establish broader generalizability. Finally, although COMPASS-derived scores integrate readily with survival analyses, they do not themselves constitute fully adjusted predictive models incorporating clinical covariates.

## Data Availability

Example expression matrices and gene-signature files used to generate Figure 2 and related analyses are available at our GitHub repository: https://github.com/sinha7290/COMPASS. The COMPASS web application implementing the composite activity scoring framework is freely accessible at https://compass.precsn.com/. Further information and requests for resources should be directed to and will be fulfilled by, Saptarshi Sinha.

## Disclosure Statement on Generative AI

Generative AI has not been used to write the manuscript or generate data figures de novo. AI-related tools were used to create schematics or to improve the readability and language of the text after it had been drafted.

## Acknowledgments

The authors wish to apologize for not citing many important publications on this topic due to the limit of references allowed. P.G. is supported by NIH grants R01-AI141630, R01-CA305983 and R01-AI55696. PG was also supported by the Leona M. and Harry B. Helmsley Charitable Trust and the Propel a Cure Foundation. S.S. was supported through The American Association of Immunologists (AAI) Intersect Fellowship Program for Computational Scientists and Immunologists.

## Author Contribution

S.S. and P.G. conceptualized the study. S.S. performed all computational analyses and developed the web tool. P.G. provided key resources. S.S. and P.G. jointly selected datasets and gene signatures, prepared figures, wrote, reviewed, and edited the manuscript. Both authors approved the final version of the manuscript.

## Online Methods

### Gene expression datasets and preprocessing

Previously published bulk gene-expression datasets used for benchmarking COMPASS are described in the main text. For each study, normalized expression matrices were obtained from the original publication, GEO/ArrayExpress, or were reprocessed from raw counts using standard RNA-seq pipelines (alignment, quantification, and normalization). When raw data were available, counts were converted to transcripts per million (TPM) or counts-per-million (CPM) with log₂ transformation. When only processed data were available (e.g., log₂ intensity values from microarrays), those matrices were used directly after confirming that sample-wise distributions were unimodal and approximately comparable across groups. Genes with zero or near-zero variance across samples were excluded because they do not contribute meaningfully to composite scores. All COMPASS analyses were performed on gene-by-sample matrices in which rows correspond to genes and columns correspond to samples, with samples annotated by user-defined biological or clinical group labels.

### Gene signatures

For all analyses in this study, gene lists and direction assignments were curated from prior experimental work, literature-derived pathways, or Boolean implication networks, as detailed in the main text.

### COMPASS composite activity scoring

Composite activity scores were calculated using COMPASS as defined in Equations (1–5) in the main text. Briefly, log₂-normalized gene expression matrices (e.g., TPM or counts-per-million) were used as input. For each gene, COMPASS estimated a data-driven activation threshold by fitting a step function to the ordered expression values and identifying the position that minimized the within-group variance, thereby classifying samples as “low” or “high” expressors. Expression deviations from this threshold were then standardized by the across-sample standard deviation to generate direction-aware z-scores. For each predefined gene set or signature, per-sample composite activity scores were obtained as the weighted average of these z-scores, with +1 weights assigned to genes expected to increase and −1 weights to genes expected to decrease with pathway activation. The resulting scores are deterministic (no permutations) and were used to generate boxplots, group-wise comparisons, and receiver operating characteristic (ROC) curves for user-specified biological or clinical contrasts. All analyses were performed using the COMPASS web application (https://compass.precsn.com).

### Survival analysis and outcome modeling

Survival analysis was performed using COMPASS-derived scores as continuous predictors. Patients’ metadata with time to event annotation were used as input. Patients were stratified into high and low groups using a *StepMiner*-derived cutoff applied to the distribution of composite scores within each cohort. Kaplan–Meier curves were used to estimate time-to-event (30-day mortality) probabilities, with significance assessed by log-rank test. Hazard ratios (HRs) and corresponding confidence intervals were estimated using Cox proportional hazards models, treating COMPASS scores as either continuous or dichotomized variables. In datasets where samples are annotated for the presence or absence of an event/outcome, as well as time to such outcome/event, the same composite metric used for classification and ROC analysis can be directly applied to survival modeling without additional transformation or retraining.

### Quantification and Statistical Analysis

For each gene module or signature, COMPASS computed a single continuous composite activity score per sample as described in the main text. These scores served as the primary quantitative readout for all downstream analyses. For group-wise comparisons, samples were assigned to user-defined biological or clinical groups, and composite scores were compared using two-sided Welch’s t-tests (unequal variances). Raw p-values were assembled into a group × group matrix and adjusted for multiple testing using both Bonferroni and Benjamini–Hochberg false discovery rate (FDR) procedures; both raw and adjusted values were exported for interpretation. For classification performance, a designated reference group was treated as the negative class and each other group as the positive class. Receiver operating characteristic (ROC) curves and corresponding areas under the curve (AUCs) were computed from composite scores using standard empirical estimators, and AUC p-values were obtained from asymptotic normal approximations. All tests were two-sided and based on complete-case data. For survival analysis log rank test, hazard ratio was also calculated. Analyses were implemented in Python within the COMPASS web application using standard scientific libraries (numpy, pandas, scipy, scikit-learn), with visualization modules generating boxplots, ROC curves, KM plot for export.

### Head-to-head comparison with GSVA and ssGSEA

Gene set activity was quantified using two widely used single-sample enrichment methods, Gene Set Variation Analysis (GSVA) and single-sample Gene Set Enrichment Analysis (ssGSEA), implemented via the GSEApy package (v1.1.13) with default parameters unless otherwise specified (including kernel type, normalization, and gene set size thresholds). GSVA estimates pathway activity by modeling the distribution of gene expression values across samples using a non-parametric kernel transformation, whereas ssGSEA computes a rank-based enrichment score independently within each sample. In parallel, a deterministic composite score was computed using the COMPASS framework. In this approach, gene expression values were transformed into standardized deviations from *StepMiner*-derived thresholds with noise margin (T*), scaled by gene-specific variance to generate dimensionless Z-scores. These normalized values were then aggregated using predefined gene-specific weights to generate a directionality-aware composite activity score at the sample level. Directionality weights (+1 for activation, −1 for repression) were assigned based on prior biological annotation of each gene and held constant across all analyses.

Method performance was evaluated by assessing the ability of GSVA, ssGSEA, and COMPASS scores to discriminate sepsis from healthy controls using ROC analysis and the corresponding AUC. Score orientation was aligned across methods such that higher values consistently corresponded to the disease (sepsis) state prior to ROC analysis. To assess robustness, stratified bootstrap resampling (n = 1000) was performed with replacement within each class using a fixed random seed to ensure reproducibility. AUC was computed for each bootstrap replicate, and the resulting distributions were used to estimate mean performance and 95% confidence intervals based on percentile statistics. Sensitivity and specificity were computed using a fixed decision threshold for each method, determined on the full, original cohort by maximizing the Youden index^32^ and applied consistently across bootstrap replicates.

Traditional GSEA was not applied because it produces cohort-level enrichment statistics rather than sample-level scores.

## REFERENCES

1. Subramanian, A. et al. Gene set enrichment analysis: a knowledge-based approach for interpreting genome-wide expression profiles. Proc Natl Acad Sci U S A 102, 15545–15550 (2005).

2. Hänzelmann, S., Castelo, R. & Guinney, J. GSVA: gene set variation analysis for microarray and RNA-seq data. BMC Bioinformatics 14, 7 (2013).

3. Barbie, D.A. et al. Systematic RNA interference reveals that oncogenic KRAS-driven cancers require TBK1. Nature 462, 108–112 (2009).

4. Tomfohr, J., Lu, J. & Kepler, T.B. Pathway level analysis of gene expression using singular value decomposition. BMC Bioinformatics 6, 225 (2005).

5. Aibar, S. et al. SCENIC: single-cell regulatory network inference and clustering. Nat Methods 14, 1083–1086 (2017).

6. Frost, H.R. Variance-adjusted Mahalanobis (VAM): a fast and accurate method for cell-specific gene set scoring. Nucleic Acids Res 48, e94 (2020).

7. Timmons, J.A., Szkop, K.J. & Gallagher, I.J. Multiple sources of bias confound functional enrichment analysis of global-omics data. Genome Biol 16, 186 (2015).

8. Wijesooriya, K., Jadaan, S.A., Perera, K.L., Kaur, T. & Ziemann, M. Urgent need for consistent standards in functional enrichment analysis. PLoS Comput Biol 18, e1009935 (2022).

9. Haynes, W.A., Tomczak, A. & Khatri, P. Gene annotation bias impedes biomedical research. Sci Rep 8, 1362 (2018).

10. Sinha, S. & Ghosh, P. (2025).

11. Sahoo, D. et al. Artificial intelligence guided discovery of a barrier-protective therapy in inflammatory bowel disease. Nat Commun 12, 4246 (2021).

12. Sahoo, D. The power of boolean implication networks. Front Physiol 3, 276 (2012).

13. Argelaguet, R. et al. MOFA+: a statistical framework for comprehensive integration of multi-modal single-cell data. Genome Biol 21, 111 (2020).

14. Argelaguet, R. et al. Multi-Omics Factor Analysis-a framework for unsupervised integration of multi-omics data sets. Mol Syst Biol 14, e8124 (2018).

15. Sahoo, D., Dill, D.L., Tibshirani, R. & Plevritis, S.K. Extracting binary signals from microarray time-course data. Nucleic Acids Res 35, 3705–3712 (2007).

16. Ghosh, P. et al. Machine learning identifies signatures of macrophage reactivity and tolerance that predict disease outcomes. EBioMedicine 94, 104719 (2023).

17. Sinha, S. et al. CANDiT: A machine learning framework for differentiation therapy in colorectal cancer. Cell Rep Med, 102421 (2025).

18. Tindle, C. et al. A living organoid biobank of patients with Crohn’s disease reveals molecular subtypes for personalized therapeutics. Cell Rep Med 5, 101748 (2024).

19. Sinha, S. et al. COVID-19 lung disease shares driver AT2 cytopathic features with Idiopathic pulmonary fibrosis. EBioMedicine 82, 104185 (2022).

20. Vo, D.T. et al. SPT6 loss permits the transdifferentiation of keratinocytes into an intestinal fate that resembles Barrett’s metaplasia. iScience 24, 103121 (2021).

21. Sinha, S., et al. F.O.R.W.A.R.D: A Data-Driven Framework for Network-Based Target Prioritization in Drug Discovery. bioRxiv (Under review at Journal of Clinical Investigation) (2025).

22. Penrose, H.M., et al. A Living Organoid Biobank of Crohn’s Disease Patients Reveals Distinct Clinical Correlates of Molecular Subtypes of Disease. *medRxiv* (2025).

23. Tindle, C. et al. Adult stem cell-derived complete lung organoid models emulate lung disease in COVID-19. Elife 10 (2021).

24. Sinha, S. et al. Growth signaling autonomy in circulating tumor cells aids metastatic seeding. PNAS Nexus 3, pgae014 (2024).

25. Sahoo, D. et al. AI-guided discovery of the invariant host response to viral pandemics. EBioMedicine 68, 103390 (2021).

26. Sweeney, T.E., Shidham, A., Wong, H.R. & Khatri, P. A comprehensive time-course-based multicohort analysis of sepsis and sterile inflammation reveals a robust diagnostic gene set. Sci Transl Med 7, 287ra271 (2015).

27. Zhang, Y. et al. A Novel Breast Cancer Index for Prediction of Distant Recurrence in HR. Clin Cancer Res 23, 7217–7224 (2017).

28. Meltzer, E.B. et al. Bayesian probit regression model for the diagnosis of pulmonary fibrosis: proof-of-principle. BMC Med Genomics 4, 70 (2011).

29. Choi, J. et al. Inflammatory Signals Induce AT2 Cell-Derived Damage-Associated Transient Progenitors that Mediate Alveolar Regeneration. Cell Stem Cell 27, 366–382.e367 (2020).

30. Liesenfeld, O. et al. Clinical validation of an AI-based blood testing device for diagnosis and prognosis of acute infection and sepsis. Nat Med 31, 4044–4054 (2025).

31. Sweeney, T.E. et al. A community approach to mortality prediction in sepsis via gene expression analysis. Nat Commun 9, 694 (2018).

32. Fluss, R., Faraggi, D. & Reiser, B. Estimation of the Youden Index and its associated cutoff point. Biom J 47, 458–472 (2005).

33. Ackermann, M., et al. Pulmonary Vascular Endothelialitis, Thrombosis, and Angiogenesis in Covid-19. N Engl J Med 383, 120–128 (2020).

34. Borok, Z. et al. Grp78 Loss in Epithelial Progenitors Reveals an Age-linked Role for Endoplasmic Reticulum Stress in Pulmonary Fibrosis. Am J Respir Crit Care Med 201, 198–211 (2020).

