## Supplemental Figures and Legends for "COMPASS enables cohort-independent digital biomarker discovery and pathway quantification"

Precision medicine · Digital biomarkers · Artificial intelligence/machine learning (AI/ML) · Transcriptomics · Cohort-independent analysis · Survival analysis · Pathway activity scoring · New Approach Methods (NAMs)· Outcome modeling

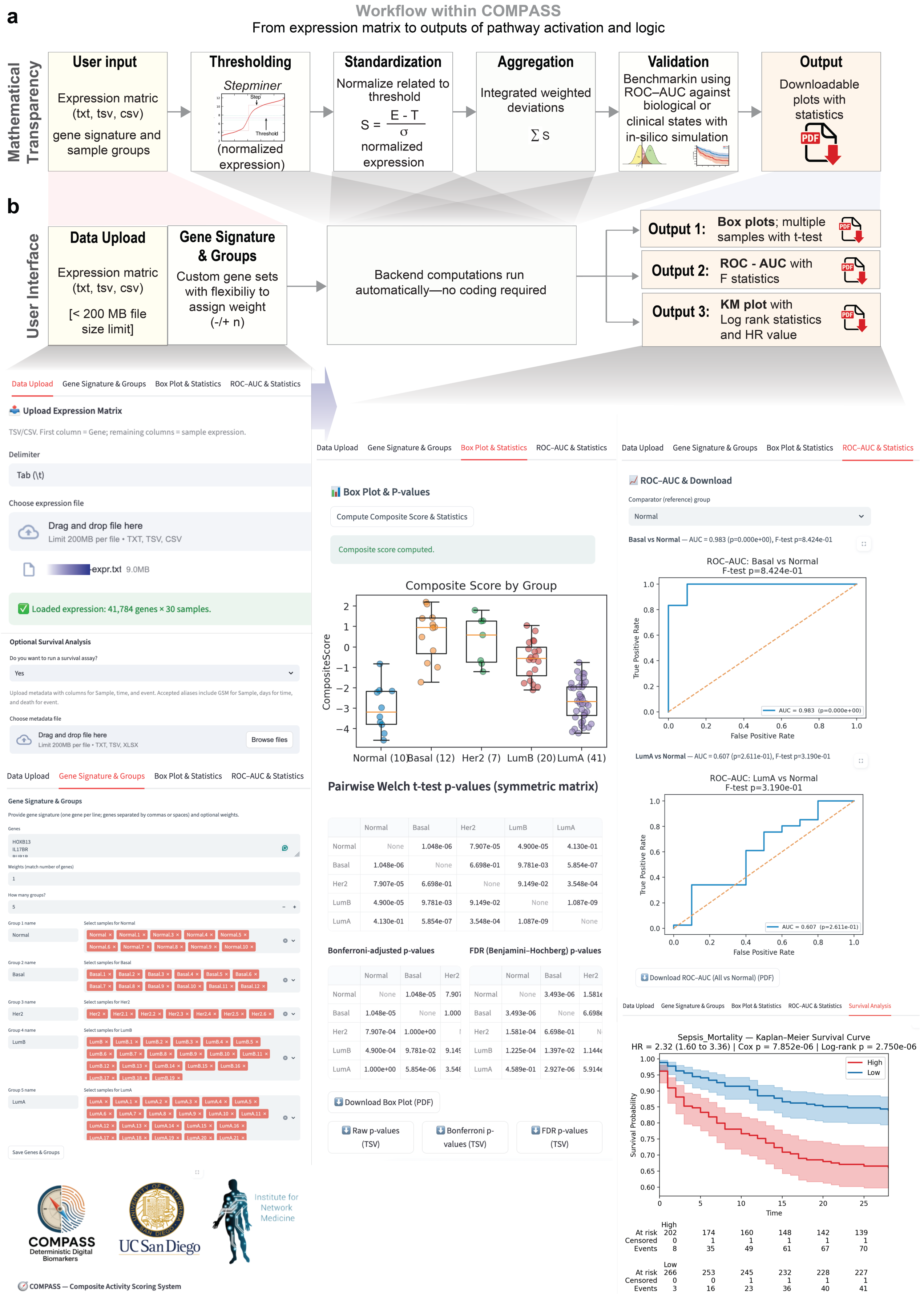


**Figure Supplement S1 [related to Figure 1].**

**Computational Workflow and User Interface of COMPASS.**

(a) **Workflow schematic**. From a user-supplied gene expression matrix (txt/tsv/csv), a sample metadata file with time to event annotation (optional), a defined gene signature (with optional weights), and sample group annotations, COMPASS executes a stepwise, transparent pipeline: (i) thresholding using *StepMiner* to define expression cutoffs, (ii) standardization of expression as deviation from threshold, (iii) aggregation of weighted deviations to generate composite pathway activity scores, and (iv) validation by benchmarking against biological or clinical states using ROC-AUC analysis and survival analysis. Outputs include publication-ready, downloadable visualizations and statistics: multi-group box plots with pairwise tests, ROC-AUC metrics with F statistics, and Kaplan–Meier curves with hazard ratio and log rank test.

(b) **User interface**. Representative screenshots illustrate the web-based interface across sequential tabs (highlighted in red): *Data Upload*, *Gene Signature & Groups*, *Box Plot & Statistics*, *ROC–AUC & Statistics*, and *Survival Analysis*. The platform enables a code-free workflow in which all backend computations are executed automatically while maintaining mathematical transparency. Example outputs are shown using clinically relevant gene expression signatures applied to human cohorts. The box plot (center) and ROC–AUC plots (right) demonstrate stratification of patients into risk groups based on PAM50 breast cancer subtypes and gene signatures used in the Breast Cancer Index® (BCI™), a clinically validated genomic assay recognized by major oncology guidelines (National Comprehensive Cancer Network [NCCN] and the American Society of Clinical Oncology [ASCO]) for guiding extended endocrine therapy decisions in HR+ early-stage breast cancer. The survival analysis (bottom right) illustrates Kaplan–Meier–based risk stratification of sepsis mortality using a 12-gene Stanford mortality signature[1], highlighting the ability of COMPASS-derived scores to associate with clinical outcomes (GSE310929).

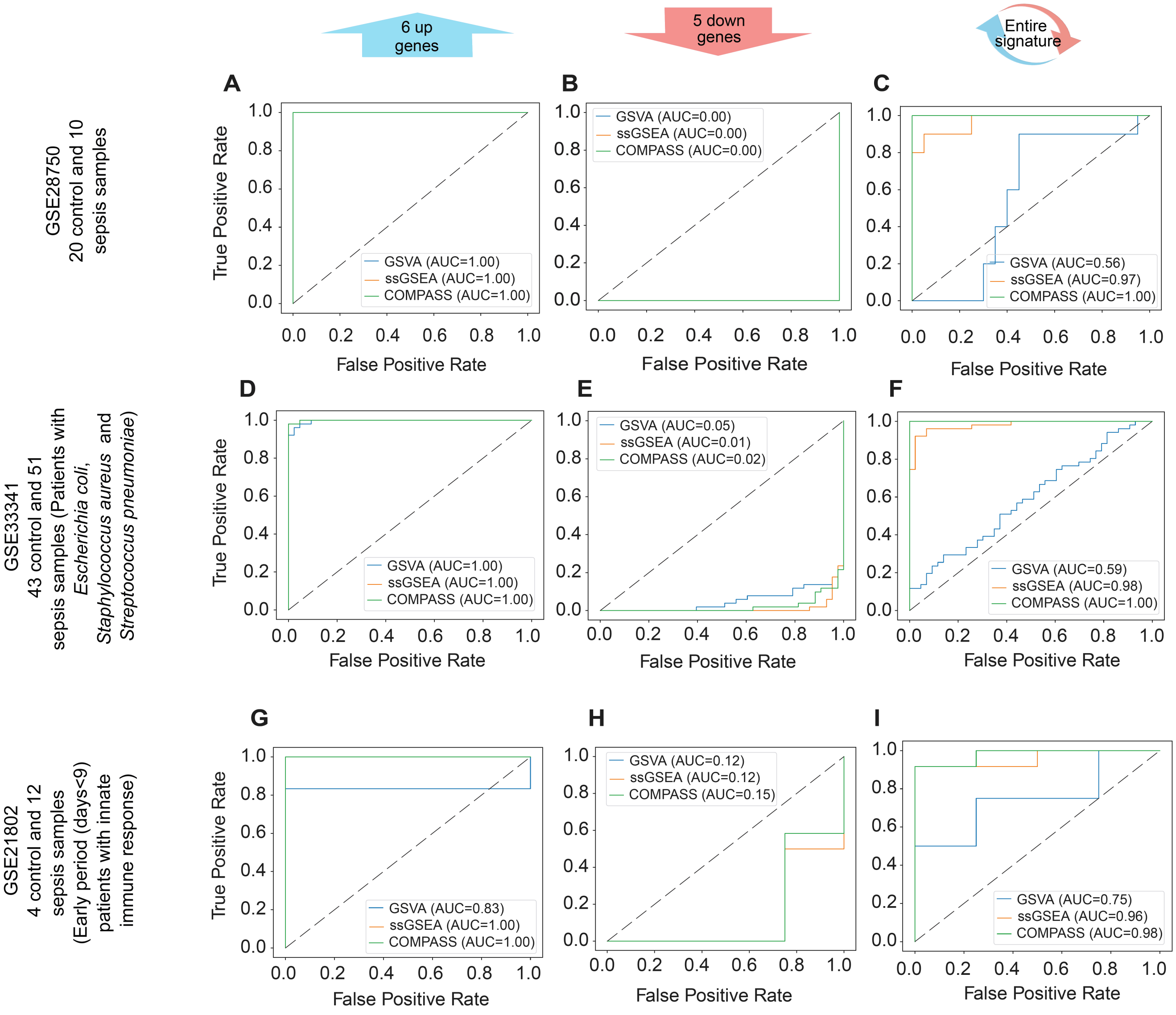


**Figure Supplement S2 [Related to Figure 3].**

**Extended multi-cohort benchmarking of COMPASS**

Receiver operating characteristic (ROC) curves comparing GSVA (blue), ssGSEA (orange), and COMPASS (green) across additional independent transcriptomic cohorts. These datasets, unique from those shown in Figure 3, represent the remaining validation cohorts for the 11-gene sepsis diagnostic signature (see **Table 2**). (**A–I**) Panels are organized by gene set composition (columns) and cohort (rows). Columns represent: Left: 6 upregulated genes, Middle: 5 downregulated genes, Right: full 11-gene signature. Rows correspond to independent cohorts: (**A-C**) GSE28750 (20 controls vs 10 sepsis), (**D-F**) GSE33341 (43 controls vs 51 bacterial sepsis; *Escherichia coli, Staphylococcus aureus,* and *Streptococcus pneumoniae*), (**G-I**) GSE21802 (4 controls vs 12 early-stage sepsis with innate immune response). Consistent with Figure 3, performance of GSVA and ssGSEA varies with gene subset and cohort context, whereas COMPASS maintains stable and high discrimination, particularly when integrating the full signature. These results extend the primary analysis and reinforce the cohort-anchored robustness of COMPASS across diverse clinical settings.

1. Sweeney, T.E., Perumal, T.M., Henao, R., Nichols, M., Howrylak, J.A., Choi, A.M., Bermejo-Martin, J.F., Almansa, R., Tamayo, E., Davenport, E.E., et al. (2018). A community approach to mortality prediction in sepsis via gene expression analysis. Nat Commun *9*, 694. 10.1038/s41467-018-03078-2.
